# A new view on functions of the lysine demalonylase activity of SIRT5

**DOI:** 10.1101/2022.11.18.517122

**Authors:** Jarmila Nahálková

## Abstract

A substrate specificity of the pharmaceutically attractive tumor-promoter SIRT5 was already investigated multiple times by advanced proteomic tools. However, the present bioinformatic analysis brings new highlights to the knowledge about the lysine demalonylation activity of SIRT5, a member of the sirtuin family with multiple roles in aging and age-related diseases. It shows unreported functional aspects of the lysine demalonylated substrates in Eukaryotic translation elongation (ETE), Amino acid and derivative metabolism (AADM), and Selenoamino acid metabolism (SAM). The cluster of the elongation factors (EEF1A1, EEF2, EEF1D, and EEF1G) belonging to ETE participates in the peptide chain elongation and the export of the tRNA-s from the nucleus to the primary sites of the proteosynthesis. SIRT5 regulates the activity of the key enzymes with tumor-promoting functions involved in AADM (GLUD1, SHMT1, ACAT1). In contrast, SIRT5 also lysine demalonylates tumor suppressor substrates as a part of the AADM and SAM interaction networks (ALDH9A1, BHMT, GNMT). It indicates comparable functions like SIRT3, which has dual tumor promoter/oncogene functions. Similar to the roles of the sirtuins, the SAM pathway impacts longevity, protects against cardiovascular diseases, and is associated with hepatic steatosis. The selen supplementation mediates the calorie restriction effect, which increases the NAD+/NADH ratio in the cells and stimulates the expression of SIRT5 and other sirtuins. SIRT5 in turn regulates the selenocysteine synthesis through the lysine demalonylation of the participating ribosomal proteins, SECISBP2 and GNMT, which creates a regulatory loop.

## Introduction

SIRT5 is a member of the NAD+-dependent sirtuin family with multiple deacylation activities. The unique character of SIRT5 within the family is its participation in reprogramming of the cellular metabolism for the metabolic demands of fast-proliferating tumor cells. Unsurprisingly, this feature attracted the scientific community’s attention to identify the main contributing factors.

SIRT5 expression is essential for cancer cell growth, such as in acute myeloid leukemia (AML) cells since the downregulation inhibits AML development in mouse models [1]. On the other hand, its upregulation during tumor cell transformation supports cancer cell proliferation [2] [3]. The genetic deletion of SIRT5 however, does not have any significant impact on the metabolism of the experimental mice except for only 60 % postnatal survival out of the expected Mendelian proportion [4].

SIRT5 possesses several enzymatic activities including weak deacetylase [5,6], but stronger desuccinylase [7] [8], deglutarylase [9], and demalonylase activities [10]. Instead, the most significant lysine deacetylation function in mitochondria is fulfilled by SIRT3 [11]. About 80 % overlap exists between the lysine acetylation and succinylation processed by SIRT5 in mitochondria [12].

According to the former studies, the succinylome processed by SIRT5 is linked to the typical mitochondrial functions, such as the tricarboxylic acid (TCA) cycle, oxidative phosphorylation, amino acid degradation, fatty acid oxidation, ketone bodies production, and glycolysis/gluconeogenesis [8] [9] [13]. Activities of succinate dehydrogenase (SDH; respiratory complex II) and pyruvate dehydrogenase (PDH) are increased in SIRT5 KO mice, and an increased rate of respiration is observed in the presence of SDH and PDH substrates. It indicates, that SIRT5 inhibits the activity of the respiratory chain complex driven through these enzymes. Previously, the desuccinylation substrate interaction network showed functions in the respiratory chain complexes I, III, and IV, glutathione-S-transferase, ribosomes, and chaperonin-containing T complex 1 (TCP1) enzymes [8]. The pathway enrichment further exhibited involvement in the functions of frataxin, and the nuclear HMGB1-HMGB2-HSC70-ERP60-GAPDH complex, which can recognize the non-natural nucleoside (thiopurine) incorporation in DNA [14]. Enriched was also the SDH-mABC1-PIC-ANT-ATPase complex containing structural and functional subunits of the mitochondrial ATP-sensitive K+ channel [8].

An interesting question can be raised about the functional differences between the specificity of the lysine succinylome, malonylome, and glutarylome regulated by SIRT5. Previously identified pathways linked to the lysine demalonylation activity of SIRT5 are not different from the desuccinylation activity since SIRT5 lysine malonylome is also associated with fatty acid degradation, glycolysis/gluconeogenesis, TCA cycle, and oxidative phosphorylation [13]. The gluconeogenesis, glycolysis, and urea cycle as the main target of SIRT5 lysine demalonylation activity was also determined by Nishida et al. (2015) [15]. The study claims that the expression of SIRT5 can increase the energy flux through glycolysis by the processing of appropriate substrates [15]. Similarly, the bioinformatics analysis of the lysine deglutarylation substrates of SIRT5 emphasized functions in the metabolism of cellular respiration, amino acids, and fatty acids metabolism [9].

On the subcellular level, the SIRT5 succinylome is mostly localized in the mitochondria, cytoplasm, and nucleus [8], while the malonylome can be typically found in the cytoplasm [15] [13] and mitochondria [15]. The cytoplasmic demalonylation should assist in increasing glycolysis/gluconeogenesis, while the mitochondrial desuccinylation serves to the intensification of oxidative phosphorylation with the increased fatty acid metabolism and diminished glycolysis/gluconeogenesis pathway [15] [13].

The present study employing the pathway enrichment and gene function prediction analysis retrieved new information about lysine malonylome regulated by SIRT5 enzymatic activity. The substrate interaction network assists in the building of a new working hypothesis about the main molecular mechanisms of the SIRT5 lysine demalonylation activity with special attention to cancer cell metabolism. It complements the current knowledge by re-using already published data and gives new ideas for further experimental research.

## Materials and methods

### The list of substrates lysine demalonylated by SIRT5

Substrates were collected from the study by Nishida et al. (2015). For the substrate selection, it was used the increased threshold with False Discovery Rate (FDR) values lower or equal to 0,01 and KO/WT (knockout/wild-type) ratio > 1,5. It yielded a substrate list of 69 proteins (Tab. 1S; Gene column 7-76), which was subjected to the pathway enrichment analysis.

### Pathway enrichment and gene function prediction analysis

Names of SIRT5 substrates not recognized directly were converted to GeneMania compatible symbols using Protein Knowledgebase Uniprot [16,17] and Online Catalogue of Human Genes and Genetic Disorders (OMIM®) [18]. GeneMania (3.5.2) analysis [19–21] was performed under Cytoscape (3.9.1) environment [22] [23] utilizing the substrate list including SIRT5 as a query (Tab. 1S, column Gene 7-76). The analysis was run against the *H. sapiens* database (update 29-04-2021) and by including all types of interaction networks. The analysis was set up for the identification of the top 20 related genes and at the most 20 attributes by employing the standard weighting method of GeneMania. It represents query gene-based weighting, which is identical to Gene Ontology (GO) Biological Processes. GO Cellular Component-based weighting also produced the main results described by the study, however with different scores. GO Molecular Function-based weighting yielded fewer results than other weighting methods (Metabolism and Metabolic pathways; Translation, and Cellular responses to stress). Scores (weights) express the predictive value, that GeneMania assigns to the pathways, and it is a measure of how well they correspond to the query dataset compared to the non-query.

### Artwork

The illustration was drawn using Adobe Illustrator Artwork 26.0 and Inkscape 1.1.1.

## Results and discussion

The tumor promoter activity of SIRT5 linked to the adaptation of the cell metabolism for the optimal growth of cancer cells is an attractive subject for tumor biologists. Determination of the targets of SIRT5 enzymatic activity leading to increased cancer cell growth might offer an alternative approach for pharmaceutical interventions.

The present study aims to provide new information about the molecular mechanisms mediating the tumor promoter activity of SIRT5. For this purpose, it was used the list of the lysine malonylated substrates mined from the study [15], which applied the label-free quantitative proteomics in the combination with the affinity enrichment by the analysis of liver lysates from WT and SIRT5 KO mice. The pathway enrichment analysis using Ingenuity Pathway Analysis (IPA) suggested that the substrates demalonylated by SIRT5 are functionally involved in the regulation of the energetic flux through glycolysis [15]. Glycolysis/gluconeogenesis are the main targets of the demalonylation activity of SIRT5 according to other authors [13].

The present analysis uses a hypothesis, that the proteins interacting directly *via* protein-protein interactions are with a high probability of participating in the same cellular and molecular functions. Mutations in the directly interacting disease genes lead to similar disease phenotypes [24]. It also uses a central model of the interaction network medicine, which relies on targeting multiple essential proteins since the treatment development for the disease pathologies through just a single node appears not feasible [25]. The most promising drug-targeting strategies are based on the partial inhibition of several protein nodes within the interaction network instead of the complete inhibition of a single node [26]. The targeting of several nodes has practical implications for the development of therapeutical approaches and the present analysis attempts to define them.

The analysis identified the general metabolic pathway, Eukaryotic translation elongation (ETE), Amino acid and derivative metabolism (AADM), and Selenoamino acid metabolism (SAM), which share protein nodes belonging to ribosomal proteins (Fig. 1, Tab. 1). The clustering and visualization of the substrate nodes assisted in the identification of tumor promoter/suppressor functions of the nodes associated with these pathways. It also revealed new information about the subcellular localization of SIRT5 lysine demalonylation activity.

**Fig. 1.**
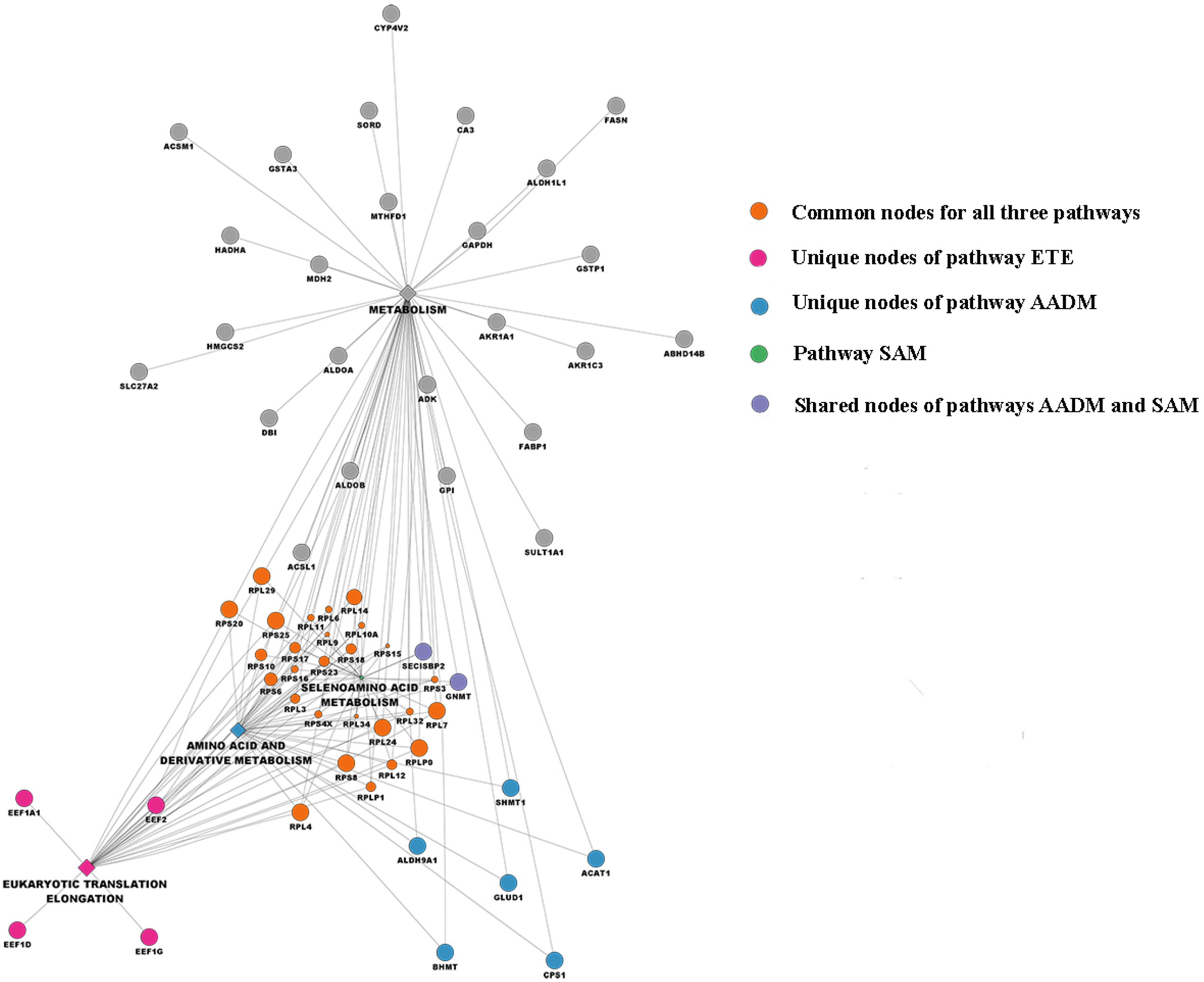
The results of the pathway enrichment and gene function prediction analysis of the interaction network of lysine malonylated substrates of SIRT5. The clusters of the most enriched pathways ETE, AADM, and SAM, and their overlapping and unique nodes are visualized in colors. The most significantly enriched general Metabolism pathway is not discussed.

**Tab. 1.**
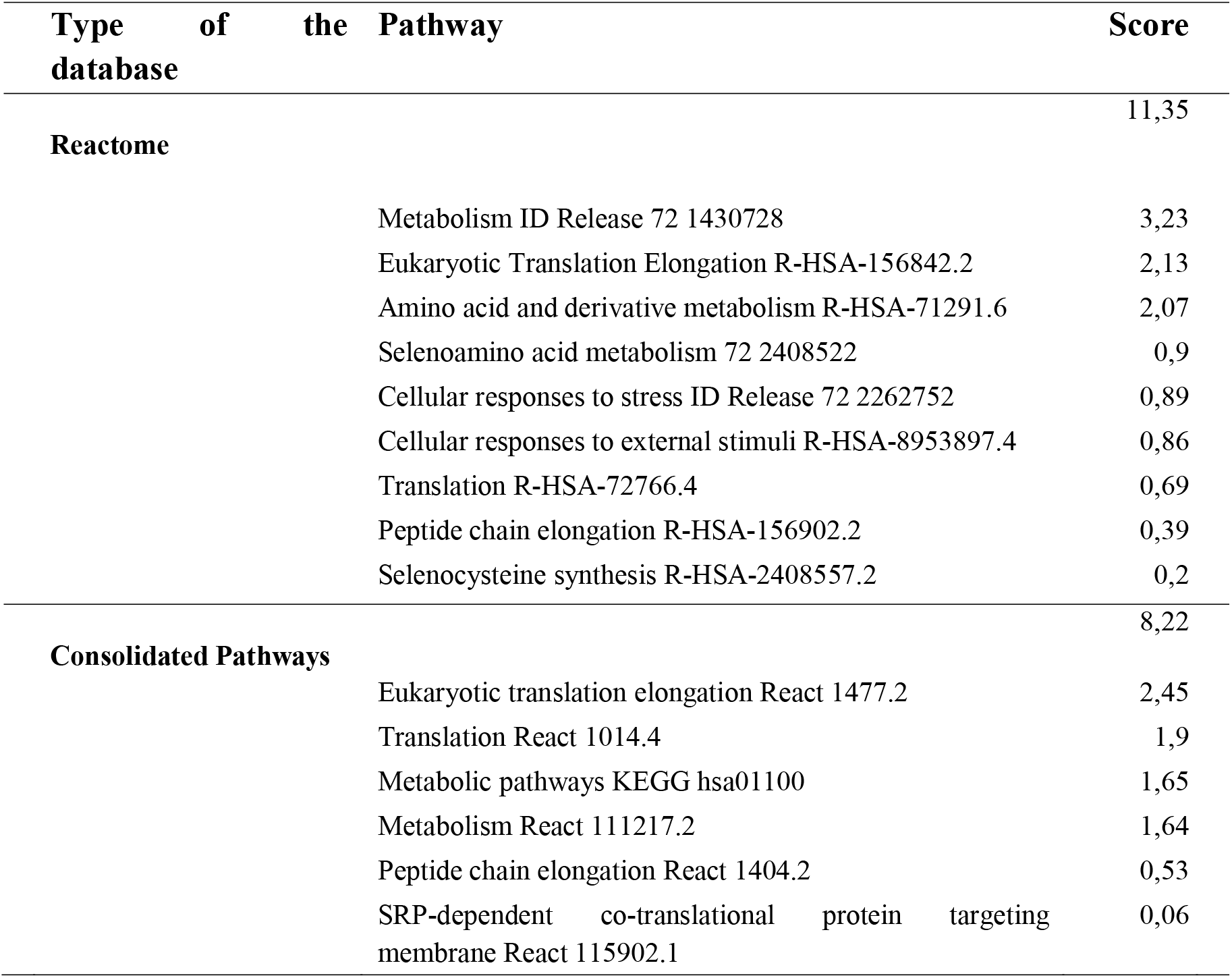
The results of the pathway enrichment and gene function prediction analysis of the interaction network of lysine malonylated substrates of SIRT5. Scores (weights) express the predictive value, that GeneMania assigns to the pathways, and how well they correspond to the query dataset compared to the non-query.

### Proteosynthesis

In addition to ribosomal proteins, the remaining substrates connected to the ETE cluster are subunits of the eukaryotic elongation factor complex (Fig. 1; EEF1A1, EEF2, EEF1D, EEF1G). Identified nodes participate in the peptide chain elongation when aminoacyl-tRNA binds to site A, delivers single amino acid to the growing polypeptide chain, and together with tRNA-s dissociates from the ribosome. A mutation in the aa-tRNA binding domain of EEF1A1 (EF1A) causes the accumulation of the tRNA in the nucleus and it proposes the function in the export of the tRNA-s from the nucleus to the primary sites of the proteosynthesis, ribosomes at the rough endoplasmic reticulum (ER) [27].

### Amino acid and derivative metabolism (AADM)

SIRT5 acts as a tumor promoter [3] [2], and the functions of its lysine malonylated substrates involved in the amino-acid metabolism support these observations.

It is known, that besides glucose metabolism, the metabolism of glycine and glutamine are the most important promoters of cancer cell growth [28] [2]. Glutamine metabolism regulated by SIRT5 is increasingly utilized in proliferating tumor cells since desuccinylation and stabilization of glutaminase (GLS), an enzyme catalyzing the conversion of glutamine to glutamate strongly promotes tumor cell growth [3].

AADM cluster also includes several enzymatic nodes with functions confirming the tumor-promoting effect of SIRT5 (Fig. 1, Tab. 1). While the function of the lysine demalonylation of GLUD1 is merely unknown, SIRT5 promotes the growth of colorectal cancer cells by deglutarylation and activation of L-glutamate dehydrogenase 1 (GLUD1; Fig. 1, Cluster AADM), an enzyme catalyzing the enzymatic conversion of glutamate into α-keto glutarate, an anaplerotic substrate of the TCA cycle. In this way, GLUD1 increases TCA flux and supports the anabolic metabolism of the cancer cells [2]. GLUD1 is also deacetylated and activated by SIRT3 [29], another member of the sirtuin family with a function in cancer, which shares multiple substrates with SIRT5.

The accelerated tumor growth is also supported by the activity of Serine hydroxymethyltransferase (SHMT1; Fig. 1, Cluster AADM), an enzyme catalyzing the conversion of serine and tetrahydrofolate to glycine, and 5, 10-methylene tetrahydrofolate [30] required for thymidine and purine synthesis [31]. The enzyme exists as cytoplasmic (SHMT1) and mitochondrial (SHMT2) isoforms [32] playing fundamental roles in cancer cell proliferation. A cytoplasmic isoform SHMT1 is a tumor promoter with a central role in the metabolic reprogramming of the cancer cells and adaptation to the tumor microenvironment [28], which fits SIRT5 functions [3]. The glycine uptake and its mitochondrial metabolism are particularly increased in the actively proliferating cancer cells. Targeting glycin dependency as a major supply of carbon e.g. by the downregulation of SHMT1 is a strategy for cancer treatment [28]. SHMT1 knockdown initiates the cell cycle arrest at the G1 phase and p53-induced apoptosis. Compared to SHMT2 downregulation, the cell cycle arrest effect is much stronger, which confirms that SHMT1 is more important for lung cancer cell survival than SHMT2 [33]. Mitochondrial isozyme SHMT2 promotes the growth of cancer cells by limiting pyruvate kinase and redirecting the glucose metabolism towards glycolysis [34]. Interestingly, SMHT2 is also acetylated, succinylated, and fatty-acylated and these modifications are removed by SIRT3 [35], SIRT5 [36], and HDAC11 [37].

Another tumor promoter present in the AADM cluster is a mitochondrial Acetyl-CoA acetyltransferase (ACAT1; Fig. 1). The physical interaction of ACAT1 with SIRT5 [38] further supports the evidence that ACAT1 is a substrate of SIRT5 [15]. SIRT5 acts as a prostate cancer promoter due to the upregulation of the MAPK pathway through ACAT1. It positively regulates the expression of ACAT1 in PC3 cells and reversely, ACAT1 positively regulates the expression of SIRT5 in LNCaP cells [38].

Contrary to the above-mentioned results, additional nodes of the AADM cluster shows tumor suppressor functions. Aldehyde dehydrogenase 9 family member A1 (ALDH9A1; Fig. 1) oxidizes a broad range of aldehyde substrates to the corresponding acids such as γ-aminobutyraldehyde to γ-aminobutyric acid [39]. Its deficiency causes DNA damage leading to genomic instability and potential tumorigenesis [40].

Another tumor suppressor node is Betaine-homocysteine S-methyltransferase (BHMT, Fig. 1, AADM), a cytosolic enzyme catalyzing the production of methionine and dimethylglycine from betaine and homocysteine [41]. It is involved in the degradation of choline to L-serine, one-carbon methylation, trans-sulphuration, cysteine, and methionine metabolism [42]. BHMT KO mice are susceptible to fatty liver and hepatocellular carcinoma, which demonstrates a protective role against hepatic steatosis and tumor suppressor functions. ALDH9A1 and BHMT are also deacetylation substrates of SIRT3 [39], which further confirms the connectivity of the SIRT3 and SIRT5 protein-protein interaction network of tumor suppressors/promoters.

A central enzyme of the ammonia detoxification pathways through the urea cycle Carbamoyl phosphate synthetase 1 (CPS1; Fig. 1; AADM), is processed by multiple SIRT5 activities. It is activated after SIRT5 deacetylation [6] [5] and deglutarylation [9] and it is a demalonylation substrate [15]. The regulation of the urea cycle through CPS1 was proven by the hyperammonemia in the blood occurring in SIRT5 KO mice accompanied by a deficiency of CPS1 activity [6]. The regulation could, however, occur through the removal of any of the mentioned three post-translational modifications and not only through its deacetylation as it was previously concluded.

The present analysis revealed that the AADM subcluster contains enzymes with tumor growth-promoting (GLUD1, SHMT1, ACAT1) and tumor suppressor activities (ALDH9A1 and BHMT). The occurrence of shared substrates with the tumor suppressor/oncogene SIRT3 proposes, that the common network of SIRT3 and SIRT5 substrates modulate the cellular microenvironment through both tumor suppressor/promoter functions.

### Selenoamino acid metabolism (SAM)

A new pathway linked to SIRT5 lysine demalonylase activity identified by the present study is the Selenoamino acid metabolism (Fig. 1, Tab. 1). Despite the functional relationships between the selen levels and the sirtuin family function were not studied in-depth [43], there exist strong indications about mutual effects.

Selenocysteine can mimic the effect of calorie restriction (CR) and extend the lifespan in nematodes [44], which fits the functions of the sirtuin family. In addition, selenocysteine decreases the high-glucose and reactive oxygen species (ROS) toxicity, which also imitates the effect of CR. Selenium supplementation regulates DNA methylation and has a protective effect on DNA, chromosome damage, telomere length, and mitochondrial DNA [45]. Further, it decreases the effect of one of the hallmarks of Alzheimer’s disease, A*β*, and the associated toxicity [44], which occurs by the inhibition of NF-κB transcription through the deacetylation of RelA/p65 [46].

Besides the functional link to CR, another common feature of the selenoprotein metabolism and SIRT5 is a connection to hepatic steatosis syndrome. Mice lacking selenocysteine lyase (ScLy), an enzyme contributing to the recycling of selenium for selenoprotein synthesis, develop a whole range of metabolic syndrome symptoms including hepatic steatosis [47]. SIRT5 deacetylates metabolism-related proteins, which results in the decrease of hepatic steatosis in ob/ob mice, an obesity mouse model. The overexpression of SIRT5 in this mouse model attenuates the hepatic steatosis symptoms [13]. It can be concluded that the mechanism of the selenium effect on hepatic steatosis is by increasing the NAD+/NADH ratio, which further stimulates SIRT5 activity [48] (Fig. 2). The results indicate that the selenoprotein metabolism and SIRT5 are linked with metabolic pathways leading to the hepatic steatosis by the lysine demalonylation of the appropriate substrates.

**Fig. 2.**
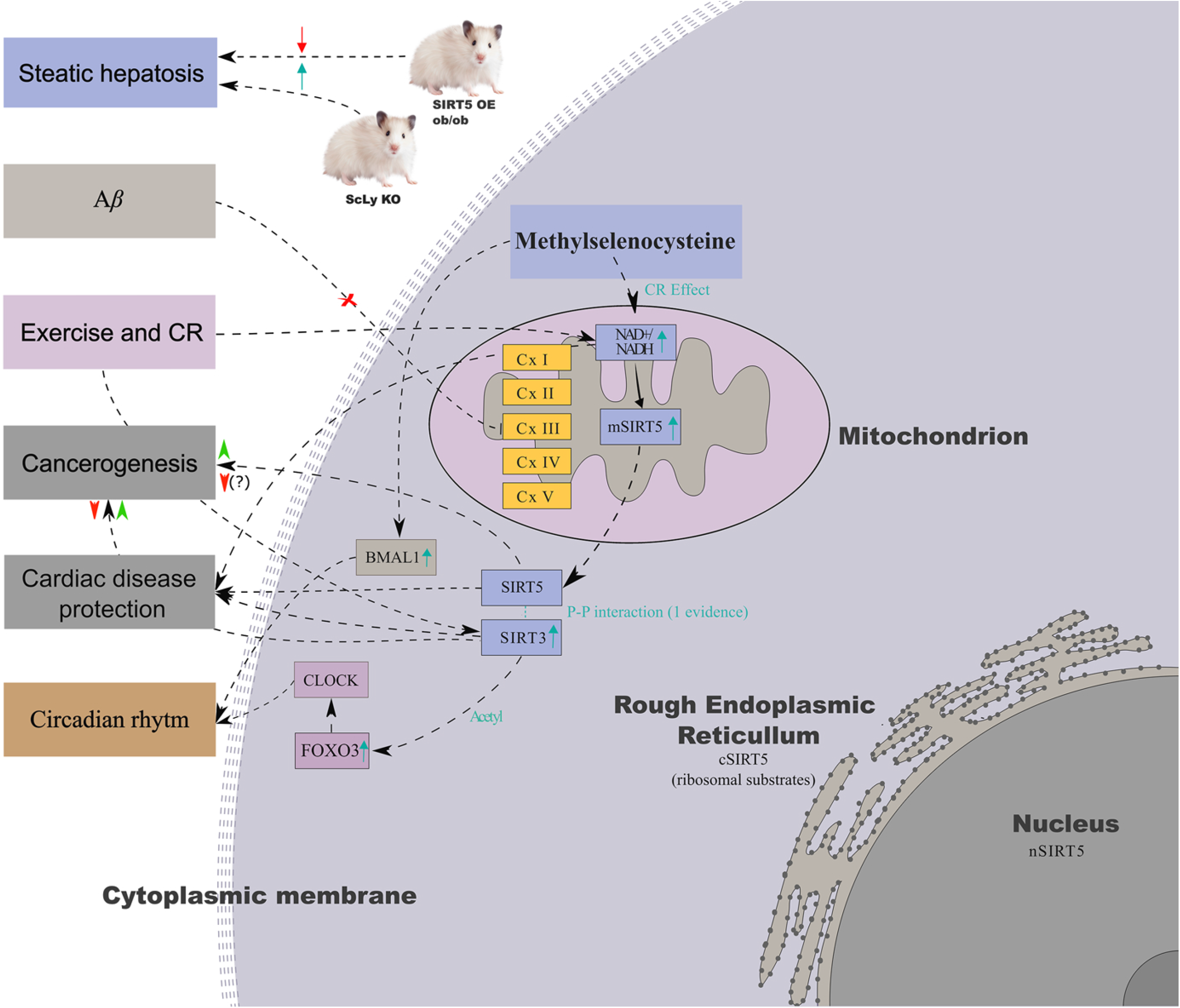
A model of the interactions between SIRT3, SIRT5, and SAM. The link between the Selenocysteine (SC) metabolism and SIRT5 functions in hepatic steatosis (HS) is shown by the example of mice lacking selenocysteine lyase (ScLy). They develop metabolic syndrome symptoms including HS, which is attenuated in the obesity ob/ob mice model by overexpression of SIRT5. SC mimics the effect of calorie restriction (CR) similar to exercise and increases the NAD+/NADH ratio, which further stimulates the activity of SIRT5 and other sirtuins. SC decreases the effect of A*β* one of the hallmarks of Alzheimer’s disease. The lysine demalonylation of the multiple substrates with tumor promoter and suppressor activities suggests dual tumor suppressor/oncogene functions similar to directly interacting SIRT3. A functional link also exists between the selenium and NAD+ levels, expression of SIRT3, SIRT5, and cardiac disease. Methylselenocystein (MSC) affects the circadian cycle by restoring the NAD^+^/NADH ratio and sirtuin activities. Stimulated SIRT1 expression deacetylates AcBMAL1, decreases its binding to the E-box element of the promoter of PER2, and rises its circadian transcription. The regulation of the circadian cycle also occurs through the SIRT3-FOXO3-CLOCK pathway, where FOXO3 is the crucial regulator. The subcellular localization is shown as mitochondrial (m), cytoplasmic (c), and nuclear (n) SIRT5.

Except for the ribosomal proteins, SAM and AADM share only two nodes, Selenocysteine insertion sequence (SECIS)-binding protein (SBP2) and glycyl N-methyltransferase (GNMT) (Fig. 1). SBP2 binds to the selenoprotein mRNA and mediates the insertion of selenocysteine into the polypeptide chain [49], and GNMT catalyzes the synthesis of N-methyl glycine from glycine by transferring methyl residue from S-adenosylmethionine. It is a tumor suppressor, which maintains genomic integrity by preventing uracil misincorporation and increasing DNA repair [50]. Colon and liver GNMT is downregulated by feeding selen-deprived feed to rat models [51], which confirms the functional link to the selen metabolism.

Methylselenocystein (MSC) also increases the expression of SIRT1, which deacetylates AcBMAL1, decreases its binding to the E-box element of the promoter of PER2, rises its circadian transcription [48], [52], (Fig. 2) and stimulates circadian expression of the genes involved in the DNA repair and suppression of cancerogenesis [48]. The regulation of the circadian cycle also occurs through the SIRT3-FOXO3-CLOCK pathway (Fig. 2) [53] [54] [55] [56] [57] where FOXO3 is a crucial regulator of CLOCK and SIRT3 is a direct SIRT5 interactor [58] [59].

Deacetylation and dessucinylation of substrates involved in mitochondrial bioenergetics are closely related to heart functions. There exists a potential link between the blood selenium levels, expression of SIRT5, and cardiac disease (Fig. 2). Interestingly, the low blood plasma levels of selenium are associated with low expression of sirtuins SIRT1, SIRT5, SIRT6, and SIRT7 and upregulation of C-reactive protein (CRP), a biomarker of the increased risk of cardiovascular disease [60] [61] [62]. SIRT5 KO mice suffer increased size of heart infarction, which is linked to the SIRT5 processed succinylome widely spread in heart tissues [63] [64]. SIRT5 reexpression in SIRT5 KO mice protects against death due to cardiac pressure overload [65]. The maintenance of NAD+ levels in the heart can prevent cardiac failure [66]. It occurs through the activation of the LKB1 by SIRT3 deacetylation, which further stimulates the LKB1-AMPK pathway [66]. SIRT3 protects the mitochondrial homeostasis against cardiomyocyte apoptosis and the development of diabetic cardiomyopathy by activating Parkin-mediated mitophagy [67]. Selenium levels, SIRT3 and SIRT5 display a common functional link to hepatic steatosis and cardiovascular disease. Selenocysteine mimics CR, impacts longevity, and protects against hepatic steatosis and cardiovascular disease. Further, it affects the expression of several sirtuins including SIRT5, which regulates the selenoamino acid metabolism and completes the functional loop.

### Subcellular localization of the lysine demalonylation substrates

The majority of the common protein nodes involved in three interaction subnetworks ETE, AADM, and SAM are ribosomal proteins (Fig. 1), which do not correspond with the generally accepted exclusively mitochondrial localization of SIRT5.

The extramitochondrial localization is supported by the occurrence of the isoform SIRT5iso1 in the cytoplasm and mitochondria of vertebral cells, while SIRT5iso2 is localized in the mitochondria of primates excluding humans [68]. Another study of SIRT5 succinylome reported a portion of SIRT5 substrates including ribosomal proteins localized in the cytoplasm of cell lines and mouse liver tissue and a minor proportion in the nucleus [8]. Processing of the histone substrates with potential impact on the chromatin functions also proposes the nuclear localization of SIRT5 [8]. The data mining shown in the present study also demonstrates the presence of histones H2BC4 and H1-0 on the substrate list (Tab. 1S) as a part of the interaction network Cellular Response to Stress and External Stimuli (Tab. 1). Low H1-0 expression is essential for tumor cell growth since its silencing allows the subset of the cancer cells to proliferate indefinitely. Re-expression suppresses the oncogenic pathways and cancer cell proliferation [69].

In conclusion, the present bioinformatic analysis suggests that substrate demalonylation occurs not only in mitochondria and cytoplasm but also in ribosomes and the nucleus. The potential regulatory effect of the lysine demalonylation of H1-0 could represent another mechanism of the tumor promoter/suppressor activity of SIRT5, which remains to be clarified.

### Interaction with SIRT3

SIRT5 with high confidence interacts with SIRT3 [58] [59], a tumor suppressor, which genetic deletion triggers a whole range of diseases including cancer, neurodegenerative, cardiovascular diseases, and metabolic syndrome [70] [53]. Recently, the dual tumor suppressor/oncogenic functions were shown to be mediated by SIRT3 [71], which suggests that both SIRT3 and SIRT5 might possess such functions.

The tumor growth-promoting/suppressive effect and the protective effect against cardiac disease and hepatic steatosis are likely mediated by their direct interaction and shared substrates. These effects raise interest to discover the differences in the substrate specificity of these two family members.

## Conclusion

The present pathway enrichment and gene function prediction analysis create ideas and a strong experimental hypothesis, which would not be feasible to create through conventional literature and data-mining methods. The results illustrate how the demalonylation of the elongation factors is involved in the peptide chain elongation and the transport of tRNA-s from the nucleus to the ribosomal location. The SIRT5 substrates promote cancer cell growth by regulating the key enzymes involved in AADM (GLUD1, SHMT1, ACAT1) and through the processing of the tumor suppressor substrates involved in AADM and SAM (ALDH9A1, BHMT, GNMT). It indicates both tumor promoter and suppressor functions based on the cellular microenvironment conditions, which should be experimentally verified.

Involvement in the selenocysteine metabolism is a new function of SIRT5, despite the effect of MSC on the increasing NAD+/NADH ratio, which mimics CR is already known. The ratio stimulates the activity of several members of the SIRT family including SIRT5, which through the lysine demalonylation of the appropriate enzymes controls the selenocysteine metabolism and creates a regulatory loop.

## Abbreviations

CR: calorie restriction
KO: knockout
NAD+: Nicotinamide Adenine Dinucleotide
NADH: Nicotinamide Adenine Dinucleotide + Hydrogen
PDH: pyruvate dehydrogenase
SDH: succinate dehydrogenase
TCA cycle: Tricarboxylic acid cycle
WT: wild type

## Acknowledgments

The research was conducted from the financial resources of Biochemworld co., Sweden.

## Consent statement/Ethical approval

Not required.

---

**Tab. S1.**
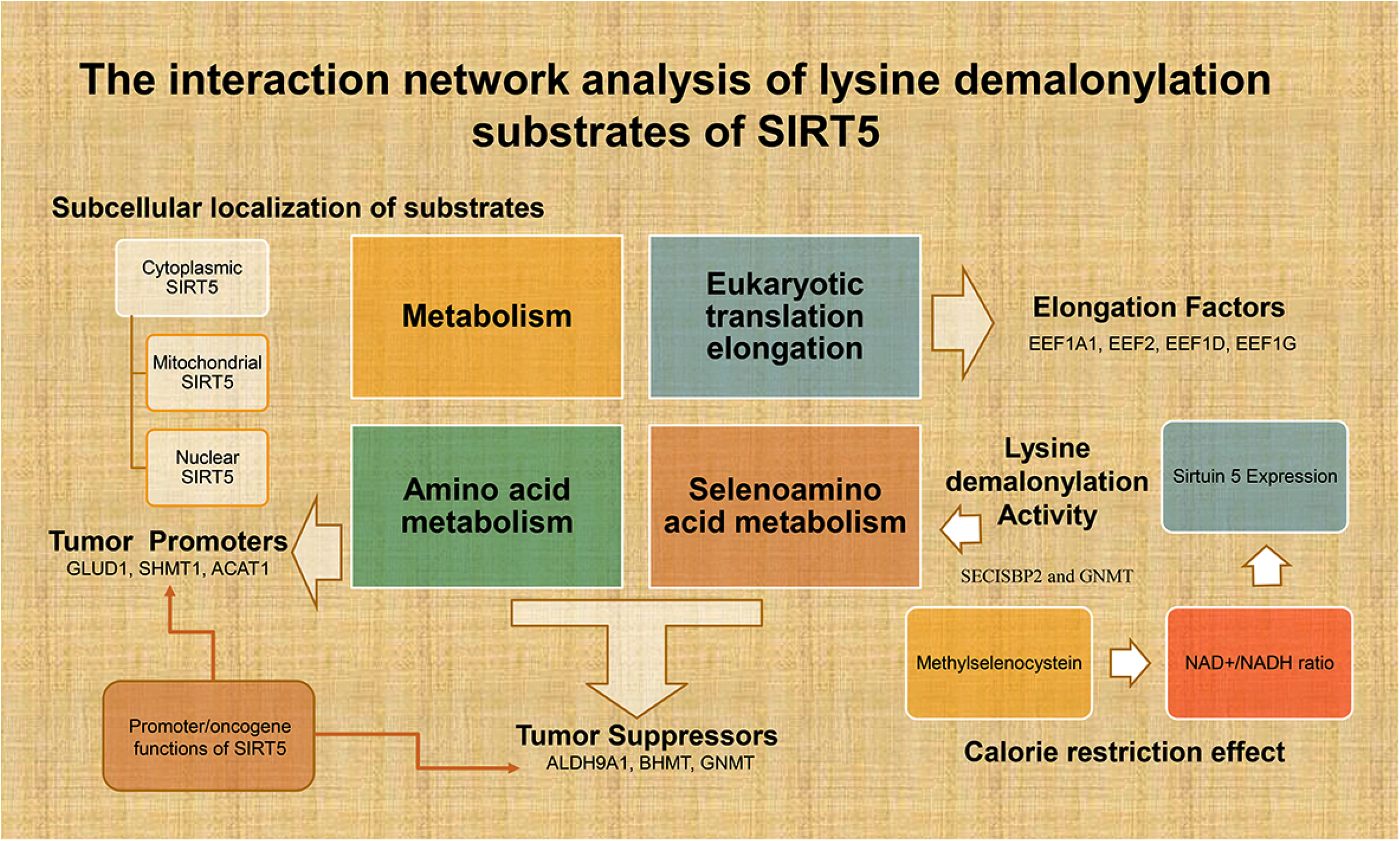
The full results of the pathway enrichment and gene function prediction analysis of the interaction network of lysine malonylated substrates of SIRT5. The table includes the input protein list (column Gene 7-76).

## Highlights

- SIRT5 lysine demalonylates elongation factors participating in proteosynthesis
- Tumor suppressor/promoter functions of lysine malonylated substrates of SIRT5
- Cancer-related functions of substrates are associated with the amino acid metabolism
- SIRT5 regulates the selenoamino acid metabolism and selenium controls SIRT5 expression

